# A Deep Learning Method for MiRNA/IsomiR Target Detection

**DOI:** 10.1101/2022.04.04.487002

**Authors:** Amlan Talukder, Wencai Zhang, Xiaoman Li, Haiyan Hu

**Affiliations:** Department of Computer Science, University of Central Florida, Orlando, FL, 32816; Burnett School of Biomedical Science, University of Central Florida, Orlando, FL, 32816; Genomics and Bioinformatics Cluster, University of Central Florida, Orlando, FL, 32816

## Abstract

**Motivation:** Accurate identification of microRNA (miRNA) targets at base-pair resolution has been an open problem for over a decade. The recent discovery of miRNA isoforms (isomiRs) adds more complexity to this problem. Despite the existence of many methods, none considers isomiRs, and their performance is still suboptimal. We hypothesize that by taking the isomiR-mRNA interaction into account and applying a deep learning model to study miRNA-mRNA interaction features, we may improve the accuracy of miRNA target predictions.

**Results:** We developed a deep learning tool called DMISO to capture the intricate features of miRNA/isomiR-mRNA interactions. Based on 10-fold cross-validation, DMISO showed high precision (95%) and recall (90%). Evaluated on three independent datasets, DMISO had superior performance to five tools, including three popular conventional tools and two recently developed deep learning-based tools. By applying two popular feature interpretation strategies, we demonstrated the importance of the miRNA regions other than their seeds and the potential contribution of the RNA-binding motifs within miRNAs/isomiRs and mRNAs to the miRNA/isomiR-mRNA interactions.

**Availability:** The source code and tool are available at http://hulab.ucf.edu/research/projects/DMISO.

**Supplementary information:** Supplementary data are available online.

## 1 Introduction

MicroRNAs (miRNAs) are ~22 nucleotides (nt) long single-stranded non-coding RNAs that play important roles in gene regulation and disease progression [1–6]. During metazoan miRNA biogenesis, miRNA genes are transcribed into pri-miRNAs, which are cut by the enzymes Drosha and DGCR8 to create the hairpin-structured pre-miRNAs. The pre-miRNAs are then exported to the cytoplasm and processed by the enzyme Dicer to produce the duplex miRNAs. Finally, the miRNAs are matured from either or both strands of the duplex miRNAs. These mature miRNAs directly bind and interact with their target mRNAs in different cell types through the context-specific choices of target sites, which results in the degradation and/or translation repression of the target mRNAs [1, 4, 6]. It is thus important to investigate how miRNAs select their target sites and identify their target mRNAs.

The discovery of different types of miRNA isoforms (isomiRs) makes the overall investigation of miRNA target sites and target genes even more challenging while fascinating [7]. During metazoan miRNA biogenesis, isomiRs are created by imprecise cleavage of pri-miRNAs and/or pre-miRNAs [8], the addition/deletion of nt to/from the ends of mature miRNAs [9, 10], and the modification of one or more nt in the middle of mature miRNAs. Correspondingly, the resulted isomiRs are classified into addition, deletion, and polymorphic isomiRs, respectively. Based on the modified end, both addition and deletion isomiRs can be further grouped as 3′ isomiRs and 5′ isomiRs. The 3′ isomiRs are more abundant and share the same seed region (positions 2-7) as their original miRNAs while the 5’ isomiRs have different seed regions and hence different target mRNAs. The resulting isomiRs can also be hybrids of the aforementioned types.

The widespread isomiRs in different cell types call for a revisit to the miRNA target site selection and target mRNA identification problem [11–14]. Under any given experimental condition, there are likely diverse changes in sequence and expression of individual miRNAs. The diversity implies the existence of different isomiRs of varied abundance and the conventional miRNAs actively interacting with their respective target genes under an experimental condition. Previous studies have shown that such a mixture of miRNAs and their isomiRs under specific experimental conditions are common instead of sequencing artifacts [15, 16]. The available methods and tools for miRNA target prediction take only the miRNAs into account, which may partially explain their high false-positive rates and sub-optimal performance [17]. It is thus critical to study isomiRs and miRNAs together for their target site and target mRNA identification.

The CLASH (crosslinking, ligation and sequencing of hybrids) and CLEAR-CLIP (covalent ligation of endogenous Argonaute-bound RNAs with cross-linking immunoprecipitation) provide an unprecedented opportunity to study miRNA targets under the context of isomiRs [18, 19]. Both experiments provide the chimeric reads composed of pairs of miRNA variants and their interacting target sites in mRNAs. These data, especially the CLASH data, have been widely used to study non-canonical target sites, which show the importance of regions other than the seed regions in miRNAs [20–25]. However, these data have not been explored to study how miRNAs and their isomiRs interact with their targets.

In this study, we attempted to predict target sites and target mRNAs for the first time by considering miRNAs together with their isomiRs. As the features of miRNA target sites and miRNA-mRNA interactions are not fully understood, and deep learning-based approaches have shown better performance in analyzing genomic and epigenomic data [26], we designed a deep learning method and tool for <u>m</u>iRNA and <u>iso</u>miRtarget target prediction (DMISO). Tested by crossvalidation and on independent datasets, we showed that, on average, DMISO had a precision of 95% and a recall of 90%. Compared with three popular tools and two recently developed deep learning-based tools, DMISO showed superior performance in almost every metric to the five tools. The DMISO tool and its codes are freely available at http://hulab.ucf.edu/research/projects/DMISO.

## 2 Materials and Methods

### 2.1 Identification of miRNA-mRNA and isomiR-mRNA interactions in CLASH

To train and test DMISO, we obtained miRNA-mRNA and isomiR-mRNA interactions from the CLASH experiments [18]. We downloaded the CLASH data in the HEK293 cell line (GSE50452), which contained raw reads from six human samples. Each sample consisted of both single and chimeric reads. Only the chimeric reads comprised the miRNA/isomiR sequences and their interacting target site sequences in mRNAs.

We identified miRNA-mRNA interactions from the chimeric reads similar to the original study [18] (Figure 1A and Supplementary Table S1). In brief, we downloaded raw reads, removed adapters from the reads, discarded duplicate reads, and finally mapped the remaining reads against two databases separately with BLAST version 2.10.1+ [27]. One database was the protein-coding transcript sequences from GENCODE version 38 [28]. The other was the human mature miRNA sequences from miRBase version 22.1 [29]. We required a BLAST hit with an evalue ≤ 0.1 to claim the mapping of a read. The mapped reads along the antisense strand of a transcript or having alignment loops were discarded. To control the mapping quality of the miRNA portion of a chimeric read, which was much shorter than the mRNA portion and thus can be harder to differentiate from the sequencing errors, we required that the same miRNA portion occurred in at least 10 chimeric reads. We allowed a maximum gap or distance of 4 nt between the mapped miRNA portion and the mapped mRNA portion in a chimeric read [18]. The miRNA and mRNA portion of a chimeric read may be mapped to multiple miRNA and mRNA transcripts, respectively. If a read was mappable to multiple miRNA or mRNA transcripts, to retain the most significant miRNA-mRNA pair, we used the following criteria in order: (1) the pair with the smaller BLAST e-values; and (2) the pair with the larger BLAST bit scores if the e-values were the same.

**Figure 1:**
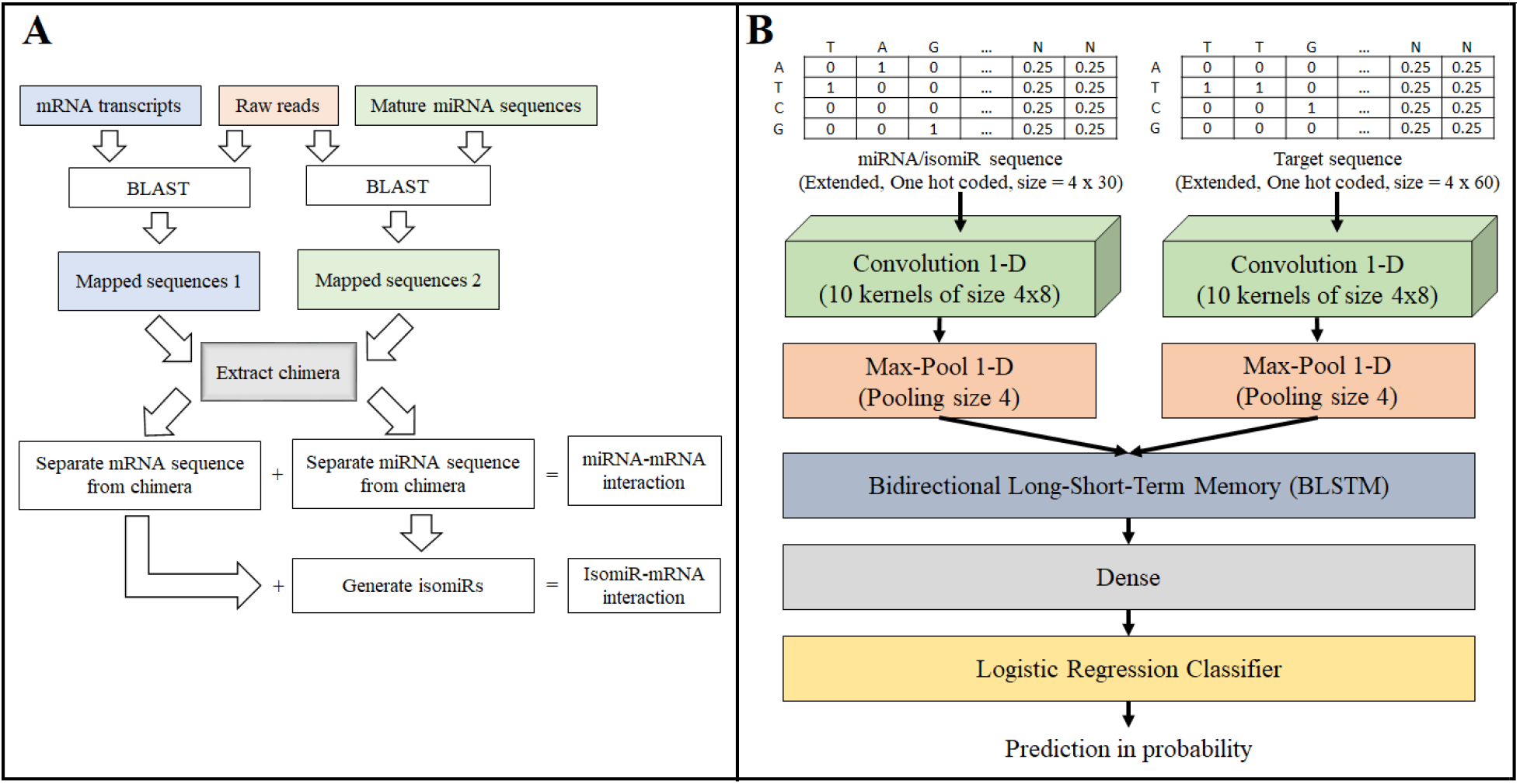
(A) The pipeline to obtain miRNA/isomiR-mRNA interactions. (B) The DMISO model structure.

With the identified miRNA-mRNA candidate pairs, we compared the aligned miRNA portion of the chimeric reads with the corresponding miRNAs to define miRNA-mRNA and isomiR-mRNA interactions. If a read perfectly matched a miRNA, we claimed this candidate pair as a miRNA-mRNA pair. Otherwise, if the nt sequencing quality scores at the variation positions (compared with the miRNA sequence) are larger than 30,this candidate pair is an isomiR-mRNA pair. To select isomiRs with confidence, we also required that the isomiR sequences were seen in at least 10 chimeric reads. We further classified these isomiRs in isomiR-mRNA pairs into the following eight types: 5’ isomiR (addition, deletion and replacement), 3’ isomiR (addition, deletion and replacement), single nucleotide polymorphic (SNP) isomiR, multiple nucleotide polymorphic (MNP) isomiR. An isomiR may belong to multiple types.

### 2.2 Training data and cross-validation

For the obtained miRNA/isomiR-mRNA pairs, we extended the 3’ end of the mRNA section of the chimeric sequences by 25 nt to have more complete target sites. The extended mRNA target sites shorter than 30 nt were filtered out as previously [18]. The extended pairs were considered as positive interaction pairs. For every positive pair, a negative pair was generated with the same miRNA or isomiR and a negative site in the 3’ untranslated region of the corresponding positive mRNA transcript. The negative site was required to be at least 10 nt away from the positive sites and had free folding energy < 10 kcal/mol measured by the RNACoFold tool [30]. With the positive and negative interacting pairs, we randomly chose 80% of positives and 80% of negatives to create a training dataset. We tested the developed DMISO method by 10-fold crossvalidation on the training dataset. We also tested DMISO on the remaining 20% of data that were not used for training.

### 2.3 Independent data

In addition to the remaining 20% CLASH test data, we extracted miRNA/isomiR-mRNA pairs from CLEAR-CLIP data as independent test data. Similar to the above analysis of the CLASH data, we analyzed the CLEAR-CLIP chimeric reads in 12 human samples from the hepatocyte-derived carcinoma cell line HuH-7.5 (GSE73059) [19]. We defined 14,684 positive miRNA/isomiR-mRNA pairs, all of them involved isomiRs instead of the conventional miRNAs.

We also obtained another independent dataset from the recently updated miRTarBase database [31]. This database contains experimentally validated functional and non-functional miRNA target sites, which are considered as positives in this dataset. We extended the 3’ end of the mRNAs in the interactions and discarded the interactions that did not have mapping mRNA and miRNA ids in the respective databases used here and those interactions where the mRNA sequences were shorter than 30 nt. After this filtering, we obtained 14,144 miRNA-mRNA sequence pairs, 13,926 of which were functional and 226 of which were non-functional interactions based on the original study [19]. This dataset did not have any negatives.

### 2.4 Deep Learning Model

We designed a deep learning method called DMISO for miRNA/isomiR target sites and target mRNA identification. DMISO takes the miRNAs/isomiRs and their corresponding mRNA target site sequences as input and outputs a binary number to indicate whether a miRNA/isomiR interacts with its corresponding mRNA site. The architecture of DMISO is composed of two separate branches containing convolutional neural network (CNN) layers, a long short-term memory (LSTM) layer, and a fully connected neural network layer (Figure 1B). The two convolutional layers are for the miRNA/isomiR and target site sequences, respectively. The LSTM layer combines the features detected by the two convolutional layers. The output of the LSTM layer is fed into a fully connected neural network to predict the label of the interaction.

The convolutional layer in each branch is 1-dimensional and consists of an array of 10 kernels, each with a size of 4 x 8. The kernels act as sliding windows to capture spatial features in input sequences by scanning the sequences. The convolutional layer does not have any padding around the input (padding = “valid”). The kernels are convolved across the input by 1 step (stride = 1). After the 10 kernels, the outputs of the two convolutional layers become the matrices of size 10 x 23 and 10 x 53, respectively. The next layer in each branch is a 1-dimensional max pooling layer with a pooling size 4, which captures the maximum values within each 10 x 4 window, sliding by 1 step (stride = 1), across the output of the respective convolutional layers. The output of the max-pooling layers in the miRNA/isomiR and target site branches is 10 x 20 and 10 x 50 matrices, respectively. Rectified Linear Unit activates the neurons in the convolutional layers of the two branches and the neurons in the dense layer. After the max-pooling step, the outputs of the two branches are merged to create a 10 x 70 matrix and fed into a bidirectional LSTM (BLSTM) layer. The BLSTM layer processes the spatially connected features from both left to right and from right to left, generating a 20 x 70 matrix output, which is then flattened to a vector of length 1400 and fed into a dense layer. The dense layer is a fully connected neural network with 100 neurons, which outputs a vector of size 100. This vector is used as an input to a logistic regression unit to generate the final prediction, where the sigmoid function is used.

Before training DMISO, the miRNA/isomiR and target site sequences are converted into 4 x 30 and 4 x 60 matrices, respectively, by applying one-hot encoding on every nucleotide in the sequences. That is, ‘A’, ‘T’, ‘C’, ‘G’ and ‘N’ are encoded into [1, 0, 0, 0]^T^, [0, 1, 0, 0]^T^, [0, 0, 1, 0]^T^, [0, 0, 0, 1]^T^ and [0.25, 0.25, 0.25, 0.25]^T^, respectively. The fixed lengths 30 and 60 are the average length of the processed miRNAs/isomiRs and target sites in chimeric reads, respectively. To keep the fixed lengths, we removed extra nt from the ends of longer sequences and added additional “N”s to the ends of shorter sequences.

Batch normalization was used to train DMISO with mini-batches of 100 samples at a time. We calculated the loss of each prediction using the binary cross-entropy loss function, which is minimized by the Adam optimizer with a learning rate of 10^-3^ [32]. To avoid overfitting, we had a dropout layer with 25% dropout rate after merging the two branches and two dropout layers with 50% dropout rate after the BLSTM layer and the dense layer. L1 regularization with the parameter value 0.01 was applied on the two convolutional layers and the dense layers to reduce overfitting. For the implementation of the deep learning model, Keras 2.3.1 version was used (https://github.com/keras-team/keras/releases/tag/2.3.1). DMISO model is executed with two inputs: miRNA/isomiR sequence and mRNA sequence The model provides an output probability score from 0 to 1 and a binary prediction value of 0 and 1.

### 2.5 Feature Identification

Many machine learning methods have been developed to select features [33–37]. Deep Learning models are infamous for being a black box when it comes to understanding the underlying features. But recent studies have focused on various strategies that can reveal the features or patterns learned by different types of machine learning models [38–42]. Here, two of the most popular feature identification methods, convolutional kernel analysis and input perturbation, were applied to discover important features for miRNA/isomiR-mRNA interactions [26].

The convolutional kernel analysis method is suitable for a deep learning model that contains a convolutional layer [26, 38–40]. This method is used to interpret the kernel weights of the convolutional layer after training the model. In this study, the miRNA/isomiR and mRNA sequences were scanned separately by the k length kernels of the two convolutional layers in DMISO, which captured the composition of k-mers in sequences that were important for the interaction between the miRNA/isomiR and mRNA sequences. Since the convolutional layers are the first layers in DMISO, the k-mer patterns captured should represent important features specific to the miRNA/isomiR and target sequences.

The input modification technique is another popular feature interpretation method [26, 38, 40], where a part of the input is perturbed with random noise and the changes in the model prediction is recorded. The change in the model prediction after the modification to a part of the input represents the sensitivity of the model to that part of the input. Therefore, this method can help reveal the model’s sensitivity patterns to different regions in input sequences. Here, we masked every contiguous region of length 4 in input sequences with “N” and recorded the respective changes in the prediction probabilities of the output layer. The changes should show important regions in terms of target binding.

### 2.6 Comparison with existing tools

DMISO was compared with three popular tools, TargetScan version 7.2 [43], miRanda 3.3 [44] and RNA22 version 2 [45], and two recently published deep learning-based tools, miRAW [46] and miTAR [47], on the 20% CLASH test data, the CLEAR-CLIP data, and the miRTarBase data. TargetScan and miRanda take two separate files for the miRNA and mRNA sequences as inputs, while miRAW and miTAR take the interactions (miRNA-mRNA sequence pairs) as inputs. In the case of isomiR-mRNA pairs in the testing data, we used the isomiR sequences in place of miRNA sequences in the input. To run RNA22, the input miRNA/isomiR and mRNA sequences must be uploaded the RNA22 server with day-wise traffic restrictions. Because of this, obtaining results from the RNA22 server on a large dataset like ours is a time-consuming process. Therefore, while the other four tools were executed on the test datasets, RNA22 was evaluated by overlapping the test data sets with the pre-computed predictions of RNA22 on human (https://cm.jefferson.edu/rna22-full-sets-of-predictions/). A test interaction was considered a predicted positive by a tool, if the miRNA id and mRNA gene id of the test interaction matched with any of the predicted interactions as well as the mRNA target sequence locations overlapped with the corresponding predicted target sites.

## 3 Results

### 3.1 The characteristics of the identified interactions in CLASH

We identified 12,170 miRNA-mRNA and 58,043 isomiR-mRNA interactions from the six CLASH samples (Supplementary Table S2). We observed each of the eight types of isomiRs, with a 3’ addition isomiR hsa-miR-4268 occurring the most frequently in 3,565 interactions, while 96 isomiRs occurring only 10 times (Supplementary Figure S1). Moreover, 98 isomiRs were found to participate in at least 100 interactions. Consistent with the previous studies [7], there were more 3’ isomiRs than other types in the CLASH data set (Supplementary Table S2). The number of isomiRs with nucleotide addition were more than other types. The number of SNP and MNP isomiRs were similar in the dataset. Despite the varied frequency of different types, there were at least 9 isomiRs from each of the eight types (Supplementary Table S3). Note that, these statistics were based on all the documented miRNAs in miRbase database. We also cross-referenced the miRNAs with curated miRNA database miRGeneDB [48] and found that 66 of the documented 268 miRNAs in miRGeneDB were in the CLASH dataset. Of them the most frequent isomiR was a 3’ addition isomiR of hsa-miR-615-3p that occurred in 824 interactions. The number of 3’ isomiRs was still higher (218) than 5’ isomiRs (57) and polymorphic isomiRs (10).

Since we considered isomiRs covered by at least 10 reads, all de novo identified isomiRs were supported by their recurrent occurrence. The mean and median number of reads supporting these isomiRs were 53 and 19, respectively. There were 200 miRNAs with at least one identified isomiR. The number of identified isomiRs for a miRNA varied from 1 to 98. On average, a miRNA had around 6 isomiRs of different types. Despite the existence of different types of isomiRs, it was evident that one miRNA preferred specific types of isomiRs. In other words, for a given miRNA, a specific type of isomiRs occurred much more frequently. In fact, for all 67 miRNAs with at least 100 isomiR-mRNA interactions, at least one type of isomiRs occurred in significantly higher than expected frequencies (corrected Binomial test p-value<0.01).

The identified isomiRs were likely to be condition-specific. We compared the identified isomiRs in the CLASH interactions with the isomiRs in the CLEAR-CLIP interactions. The CLASH samples were from a healthy kidney cell line, while the CLEAR-CLIP samples were from a carcinomic liver cell line. Among the 1,226 isomiRs and exact miRNAs identified in the CLASH data, 1,203 (98.12%) were not identified in the CLEAR-CLIP data. If we considered the 1,095 isomiRs and exact miRNAs supported by at least 50 reads in the CLASH data, 1,078 (98.45%) were still not identified in the CLEAR-CLIP data. It was thus not the isomiR quality that made the difference of isomiRs in different experiments. In other words, isomiRs and their interacting sites are likely to be condition-specific.

We investigated the difference between the unique target sites of the 5’and 3’isomiRs and those of the exact miRNAs in the CLASH dataset. Among the 5,742 CLASH target sites of exact miRNAs, 355 were common with the 5’ isomiRs’ while 2,353 were common with the 3’ isomiRs’. The lower number of common targets of exact miRNAs with the 5’ than the 3’ isomiRs corroborates that the 5’ isomiRs have altered target specificity. Interestingly, the 3’ isomiRs targeted much more unique mRNAs (5,021) than other types, indicating that the 3’ miRNA regions may also be important for miRNA/isomiR targeting.

### 3.2 DMISO had good performance on miRNA and isomiR target prediction

We evaluated DMISO by 10-fold cross-validation on the training data (Supplementary Table S4). It showed over 99% area under the receiver operating characteristic curve (AUROC) and area under the precision-recall curve (AUPR), and more than 93% F1 scores, precision, and recall.

We also evaluated DMISO on three independent datasets: the 20% left-out CLASH test data, the CLEAR-CLIP data, and the miRTarBase data (Table 1). The performance of DMISO on the left-out CLASH test data was similar to that of the above cross-validation on the CLASH training data. That is, the AUROC and AUPR were over 99%, and the F1 score, precision and recall were more than 93%. When DMISO was tested on the CLEAR-CLIP data, the performance was slightly lower (94% AUROC, 99% AUPR, 94% F1, 98% precision and 90% recall). Because we did not have negative pairs in the miRTarBase dataset, we could only evaluate the recall of DMISO on this independent dataset (Table 2). DMISO had a recall of 92%, almost as good as the recall on the CLEAR-CLIP and 20% left-out CLASH test data.

**Table 1:** Performance comparison on CLASH 20% test data and CLEAR-CLIP.

| Data set | Tools | Pos | Neg | AUROC | AUPR | F1 | Precision | Recall | Specificity |
| --- | --- | --- | --- | --- | --- | --- | --- | --- | --- |
| CLASH test | DMISO | 14053 | 8319 | 0.9945 | 0.9969 | 0.9581 | 0.9286 | 0.9895 | 0.8714 |
|  | miRanda | 14053 | 8319 | 0.5873 | 0.6921 | 0.3032 | 0.9851 | 0.1792 | 0.9954 |
|  | RNA22 | 14053 | 8319 | 0.5002 | 0.6283 | 0.0023 | 0.7273 | 0.0011 | 0.9993 |
|  | TargetScan | 14053 | 8319 | 0.5568 | 0.6686 | 0.2181 | 0.9563 | 0.1231 | 0.9905 |
|  | miRAW | 14053 | 8319 | 0.6119 | 0.6889 | 0.6552 | 0.7305 | 0.5940 | 0.6298 |
|  | miTAR | 14053 | 8319 | 0.5970 | 0.6832 | 0.5300 | 0.7639 | 0.4057 | 0.7882 |
| CLEAR-CLIP | DMISO | 14684 | 1323 | 0.9427 | 0.9945 | 0.9398 | 0.9844 | 0.8990 | 0.8420 |
|  | miRanda | 14684 | 1323 | 0.5230 | 0.9211 | 0.1135 | 0.9790 | 0.0603 | 0.9856 |
|  | RNA22 | 14684 | 1323 | 0.4993 | 0.9173 | 0.0004 | 0.6000 | 0.0002 | 0.9985 |
|  | TargetScan | 14684 | 1323 | 0.6001 | 0.9337 | 0.3670 | 0.9901 | 0.2252 | 0.9751 |
|  | miRAW | 14684 | 1323 | 0.5704 | 0.9283 | 0.7030 | 0.9367 | 0.5626 | 0.5782 |
|  | miTAR | 14684 | 1323 | 0.6904 | 0.9482 | 0.6596 | 0.9793 | 0.4972 | 0.8836 |

**Table 2:** Performance comparison on miRTarBase data.

| Tools | Recall (Overall) | Recall (Functional) | Recall (Non-functional) |
| --- | --- | --- | --- |
| DMISO | 0.9164 | 0.9168 | 0.8938 |
| miRanda | 0.7045 | 0.7054 | 0.3540 |
| RNA22 | 0.0199 | 0.0195 | 0.0221 |
| TargetScan | 0.7632 | 0.7643 | 0.3053 |
| miRAW | 0.6734 | 0.6696 | 0.5531 |
| miTAR | 0.0090 | 0.0090 | 0.0044 |

The above analysis was on all interactions in the three test datasets. We further studied how well DMISO predicted the interactions involving different isomiR types instead of exact or wild-type miRNAs (Supplementary Table S5). DMISO showed consistently good performances on different types of isomiR-mRNA interactions. For instance, DMISO had an AUROC of 94%, an AUPR of 99%, a F1 score of 93%, a precision of 98%, and a recall of 89% on the CLEAR-CLIP data for the 5’ isomiR-mRNA interactions. Overall, although the performance was close for different types of isomiR-mRNA interactions, it was slightly better on 3’ isomiR-mRNA interactions on both CLASH and CLEAR-CLIP datasets.

We also compared DMISO with five other tools: miRanda, RNA22, TargetScan, miRAW and miTAR. The input to every tool was the pairs of positive or negative miRNA/isomiR-mRNA target sites. DMISO showed a superior performance to all five tools in terms of every metric considered (Tables 1 and 2). For instance, DMISO had an AUROC of 94% and an AUPR of 99% on the CLEAR-CLIP data, compared with the highest AUROC of 69% and the highest AUPR of 95% by the other five tools. The five existing tools had a smaller precision and a much smaller recall than DMISO, which may be because DMISO was the first tool that took the isomiR-mRNA interactions into account to train the models. It also highlighted the importance of considering such interactions for future miRNA target predictions.

Among the other five tools, the deep learning-based miRAW and miTAR had slightly higher AUROC and AUPR than those of the three classical tools (TargetScan, miRanda and RNA22) and much higher recall scores. This indicates that miRAW was able to capture the information in the non-seed regions, unlike the three tools that focused on the seed regions. This implied the importance of the non-seed regions for miRNA target site identification [23, 24, 49]. On the miRTarBase dataset, DMISO had a recall that was at least 10% larger than the recall scores of other tools. In contrast, miRAW and miTar had a worse recall than miRanda and TargetScan, suggesting that the deep learning models used in the two tools might not consider certain well-studied features of miRNA target interaction captured by the DMISO model (Table 2).

### 3.3 Features analyzed

Despite the high accuracy of deep learning models to solve a problem, these models are infamous for being unable to interpret what they learn from the data. To address this problem, we used two feature interpretation methods, convolutional kernel analysis and input perturbation [26], as an attempt to unwind the learning process of the DMISO model.

Since DMISO has convolutional layers, we applied the convolutional kernel analysis method to extract the higher-level features learned by the model. In this process, the 10 kernel matrices, each of size 4 x 8, of the two convolutional layers for the miRNA (isomiR) and target branches were analyzed to find the weights of the A, T, C and G learned by the layers. In the beginning of the training process, the weight values of the kernels were randomly initialized in the forward pass and updated through backpropagation of the loss. After 500 epochs of training, 3 of the 10 kernel matrices in the miRNA (isomiR) branch and 7 of the 10 kernel matrices in the mRNA branch picked up certain weight values that might be commensurate with the protein binding motifs in the miRNA and mRNA sequences. For the remaining kernels in the respective branches, the weight values were too similar (25%) to be assigned to one of the 4 nucleotides. To find out the binding protein motifs, we compared the 4 x 8 kernel matrices of both the miRNA and target branches with the JASPAR motif database on vertebrates [50]. The top motifs that each kernel matched with high significance were the protein binding motifs GATA1::TAL1, ZFP42, RARA::RXRG, RARA::RXRA, ESR2, ZFP42, ZBTB26, etc., all of which are CCHH and CCCH type Zinc finger proteins. Zinc finger proteins are well known as RNA binding proteins, which are essential to bind with ribonucleoproteins in a RNA-induced silencing complex [51]. This analysis shows that DMISO was able to recognize the binding profile of RNA binding motifs in miRNA/isomiR and mRNA sequences through the kernels of the convolutional layers, which has not been considered in the existing miRNA target prediction tools.

The other feature interpretation method that we used was the input perturbation technique. In this case, for every miRNA and target sequence in a dataset, a 4-nucleotide long mask consisting of “N” was applied on the sequence starting at every position of the miRNA and the target sequences. The changed value of DMISO’s prediction probability after applying the mask was recorded for the region in the corresponding sequences. The mask was then slid across the miRNA and the target sequences by one nucleotide every time. The average changes in the prediction probability after scanning all the miRNA and target sequences within a dataset gave us the regions that were most significant to DMISO within all the miRNA and target sequences.

The input perturbation method was applied to the 20% CLASH test data and CLEAR-CLIP datasets (Figure 2). For each position, the mean (middle blue) and variation (gray area) of the sensitivity of DMISO to the changes at that position for all miRNA/isomiR and mRNA sequences in a dataset was recorded. In both datasets, positions 1-9 of the miRNAs/isomiRs had the most variation, which confirmed the importance of the seed region. However, not all positions in the seed were of the same importance. For instance, the first position was associated with the lowest variation while positions 3-5 were the highest in the seed for both datasets. The sensitivity variation of DMISO was also similar for both datasets from position 1 until position 18, and then dramatically decreased to the end. This might be because of the difference in miRNA/isomiR lengths (the shortest miRNA/isomiR was 17 nt long). It also suggested the importance of almost all miRNA positions instead of only the seed for target binding. When the target mRNA sequences were changed, DMISO reacted more on the modifications in the 5’ regions of the target sequences, which further implied the importance of the 3’ regions of miRNAs/isomiRs that bound the 5’ region of the target mRNA sequences. In addition to the high sensitivity variation in the 5’ region, the positions around 50 (3’ region) of the target sequences showed a spike in sensitivity variation in both datasets. Since the 3’ region of the target corresponded to the seed region of the miRNA/isomiR, this confirmed the importance of the high-quality match in this seed region as well.

**Figure 2:**
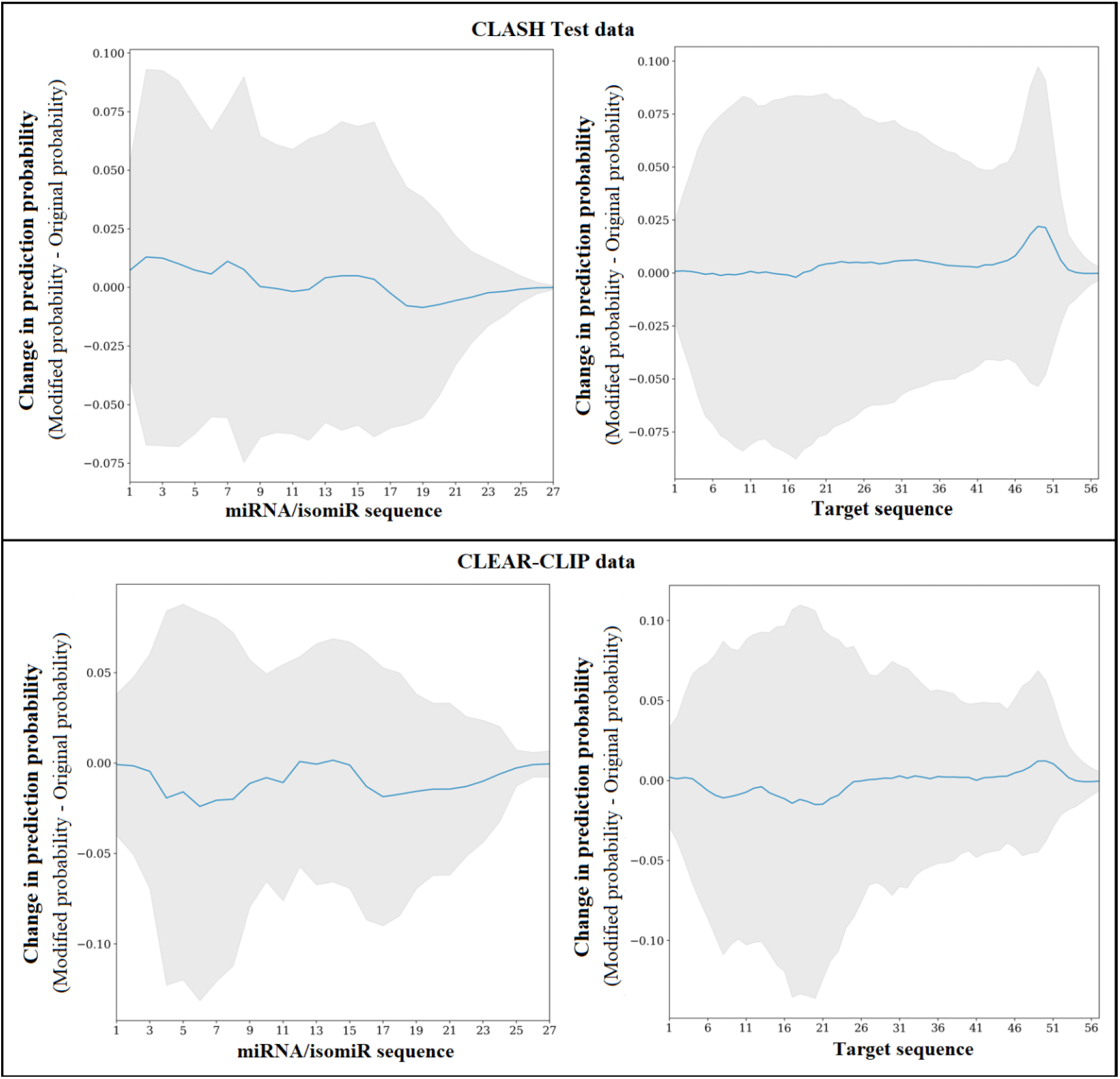
The changes in DMISO’s prediction probabilities with modification to different regions of the input miRNA (isomiR) and target sequences.

We also clustered miRNAs based on the binding sensitivity scores at every position of miRNAs. Our intuition was, if the model showed similar reaction patterns to the changes to two miRNAs, the two miRNAs might have similar binding patterns or features (Supplementary Figure S2). We found that most of these miRNA clusters had many common targets, and a cluster of miRNAs often work in the same pathways (Supplementary Table S6).

## 4 Discussion

We developed a new method DMISO to efficiently predict miRNA/isomiR targets from mere sequence inputs. The abundance of IsomiRs in different cell lines and cell types makes it impossible to ignore for miRNA target prediction. The consideration of isomiRs in DMISO enables us to consider the intricate sequence changes that contribute to the miRNA/isomiR-miRNA interactions and thus make a more informed decision in identifing miRNA target sites and targets. DMISO showed high performance on the cross-validation and three independent datasets. It outperformed the existing tools on these datasets, including three popular tools and two recently published deep learning-based tools.

We applied two feature interpretation methods to understand miRNA/isomiR-mRNA target site interaction features. The convolutional kernel analysis suggested the role of the RNA-binding protein-specific regions in miRNAs/isomiRs and the target mRNAs to form an interaction. The input perturbation technique confirmed that the 5’ region of the miRNA/isomiR and the 3’ region of the target sequence were highly important for their interactions. Additionally, it showed that the middle part of the miRNA (isomiRs) and the target sequence might also have significant contributions.

We also analyzed the performance of the model after incorporating the abundance of miRNAs/isomiRs and mRNA transcripts. The read coverage information of miRNAs/isomiRs and mRNAs were added to the inputs of the final logistic regression layer. After training the model with 500 epochs, the model showed 99% and 94% AUROC, 100% and 99% AUPR scores on the CLASH test and CLEAR-CLIP datasets, respectively, which slightly increased compared with the original DMISO model on the respective datasets. This analysis indicates that the abundance of information can improve the model. Since it is not straightforward to obtain the abundance of information in practice, we preferred the original trained DMISO model.

We did not use available tools to detect isomiR in chimeric reads because such a tool took away the flexibility to define a specific length of sequence change in an isomiR. In addition to considering only the top one pair of miRNA/isomiR-mRNA target sites from a chimeric read above, we also tried the top five potential pairs from each chimeric read to generate training and testing data. The conclusions made above still held, especially the performance of DMISO (Supplementary Table S7). With a better understanding of isomiR-mRNA interactions, we may convert chimeric reads to miRNA/isomiR-mRNA target sites better in the future.

## Supporting information

Supplementary result

## Authors’ contributions

HH and XL conceived and designed the study. AT performed the experiments. AT, XL, and HH analyzed the data. AT, XL and HH wrote the manuscript.

