## Supplementary result for "A Deep Learning Method for MiRNA/IsomiR Target Detection"

| **Table S1:** Samples, reads and aligned chimeric reads for CLASH and CLEAR-CLIP data | | | | | |
| --- | --- | --- | --- | --- | --- |
|  | Sample | Run | # Reads | # Aligned Chimeric Reads  (Alignments in minus strand and with gaps filtered) | # Aligned Chimeric Reads  (Alignment pairs with more than 4 nt gap or overlap filtered) |
| CLASH | E1_A12 | SRR959751.2 | 5304756 | 64603 | 9821 |
|  | E2_0727 | SRR959752.2 | 1194099 | 34454 | 6063 |
|  | E2_0727HS | SRR959753.1 | 9398823 | 78246 | 11115 |
|  | E3_L1 | SRR959754.2 | 1304661 | 45637 | 11201 |
|  | E3_L1HS | SRR959755.1 | 7216467 | 77193 | 20359 |
|  | E4_L2 | SRR959756.2 | 2598477 | 100448 | 24793 |
|  | E4_L2HS | SRR959757.1 | 16428827 | 213935 | 68545 |
|  | E5_TA1 | SRR959758.2 | 6629002 | 86710 | 11174 |
|  | E6_TAON | SRR959759.2 | 5600597 | 70655 | 10360 |
| CLEAR-CLIP | 1 | SRR2413175.1 | 8795072 | 42596 | 7672 |
|  | 2 | SRR2413176.1 | 2384028 | 12303 | 2963 |
|  | 3 | SRR2413177.1 | 4686830 | 7750 | 676 |
|  | 4 | SRR2413178.1 | 6794961 | 33624 | 7540 |
|  | 5 | SRR2413179.1 | 1699691 | 25244 | 10836 |
|  | 6 | SRR2413180.1 | 977323 | 13738 | 4184 |
|  | 7 | SRR2413181.1 | 1656605 | 17950 | 3272 |
|  | 8 | SRR2413182.1 | 859801 | 14069 | 2277 |
|  | 9 | SRR2413183.1 | 2619165 | 34505 | 5272 |
|  | 10 | SRR2413184.1 | 1552306 | 17394 | 2303 |
|  | 11 | SRR2413185.1 | 1433937 | 21114 | 3647 |
|  | 12 | SRR2413186.1 | 1097284 | 14426 | 2396 |

**Table S2:** Number of exact miRNAs, different types of isomiRs, targets, interactions and reads in the positive and negative interactions of CLASH training, test, CLEAR-CLIP and miRTarBase datasets. The number inside the parenthesis represents the number of miRNAs that produced the corresponding isomiRs.

|  | EXACT | 5' | 3' | Polymorphic | # Total isomiRs | # Targets | # Interactions | # Reads |
| --- | --- | --- | --- | --- | --- | --- | --- | --- |
| CLASH (pos) | 122 | 447 (99) | 957 (188) | 281 (61) | 1104 (217) | 10415 | 70213 | 70213 |
| CLASH (neg) | 120 | 332 (78) | 814 (169) | 192 (47) | 931 (197) | 5254 | 41646 | NA |
| CLASH Training (pos) | 122 | 447 (99) | 957 (188) | 281 (61) | 1104 (217) | 9713 | 56160 | 56160 |
| CLASH Training (neg) | 119 | 325 (77) | 800 (166) | 185 (46) | 914 (196) | 4988 | 33327 | NA |
| CLASH Test (pos) | 121 | 423 (93) | 921 (186) | 265 (57) | 1056 (213) | 5004 | 14053 | 14053 |
| CLASH Test (neg) | 113 | 263 (66) | 657 (144) | 147 (36) | 751 (177) | 3029 | 8319 | NA |
| CLEAR-CLIP (pos) | 0 | 691 (161) | 429 (142) | 205 (64) | 764 (170) | 975 | 14684 | 14684 |
| CLEAR-CLIP (neg) | 0 | 270 (92) | 180 (79) | 108 (39) | 295 (99) | 360 | 1323 | NA |
| mirTarBase | 573 | NA | NA | NA | NA | 5277 | 14144 | NA |

**Table S3:** Number of eight types of isomiRs occurring in CLASH and CLEAR-CLIP datasets. The isomiRs representing multiple types are shown under “Hybrid” column.

|  | 5' add | 5' del | 5' rep | 3' add | 3' del | 3' rep | SNP | MNP | Hybrid |
| --- | --- | --- | --- | --- | --- | --- | --- | --- | --- |
| CLASH | 20 | 39 | 9 | 361 | 96 | 92 | 12 | 11 | 464 |
| CLEAR-CLIP | 259 | 0 | 14 | 14 | 1 | 7 | 0 | 0 | 469 |

| **Table S4:** 10-fold cross validation on the 80% training data | | | | | | | | | |
| --- | --- | --- | --- | --- | --- | --- | --- | --- | --- |
| Pos | Neg | AUROC | AUPR | F1 | MCC | Accuracy | Precision | Recall | Specificity |
| 5616 | 3333 | 0.9952 | 0.9973 | 0.9592 | 0.8880 | 0.9473 | 0.9311 | 0.9891 | 0.8767 |
| 5616 | 3333 | 0.9952 | 0.9972 | 0.9639 | 0.9012 | 0.9534 | 0.9373 | 0.9922 | 0.8881 |
| 5616 | 3333 | 0.9933 | 0.9963 | 0.9574 | 0.8828 | 0.9449 | 0.9297 | 0.9868 | 0.8743 |
| 5616 | 3333 | 0.9950 | 0.9971 | 0.9589 | 0.8871 | 0.9466 | 0.9275 | 0.9925 | 0.8692 |
| 5616 | 3333 | 0.9938 | 0.9966 | 0.9588 | 0.8868 | 0.9468 | 0.9321 | 0.9872 | 0.8788 |
| 5616 | 3333 | 0.9945 | 0.9968 | 0.9590 | 0.8874 | 0.9468 | 0.9285 | 0.9916 | 0.8713 |
| 5616 | 3333 | 0.9951 | 0.9973 | 0.9617 | 0.8948 | 0.9504 | 0.9335 | 0.9916 | 0.8809 |
| 5616 | 3332 | 0.9947 | 0.9970 | 0.9599 | 0.8899 | 0.9481 | 0.9316 | 0.9900 | 0.8776 |
| 5616 | 3332 | 0.9939 | 0.9966 | 0.9555 | 0.8774 | 0.9422 | 0.9246 | 0.9886 | 0.8640 |
| 5616 | 3332 | 0.9941 | 0.9967 | 0.9601 | 0.8903 | 0.9485 | 0.9340 | 0.9877 | 0.8824 |

**Table S5:** Performance comparison on the interactions involving different types of isomiRs in the CLASH 20% test data and CLEAR-CLIP data.

|  | IsomiR Types | Pos | Neg | AUROC | AUPR | F1 | MCC | Accuracy | Precision | Recall | Specificity |
| --- | --- | --- | --- | --- | --- | --- | --- | --- | --- | --- | --- |
| CLASH test | EXACT | 2414 | 2065 | 0.9847 | 0.9879 | 0.9133 | 0.8057 | 0.9002 | 0.8591 | 0.9747 | 0.8131 |
|  | 5' | 3789 | 1086 | 0.9960 | 0.9989 | 0.9732 | 0.8745 | 0.9573 | 0.9520 | 0.9952 | 0.8250 |
|  | 3' | 9603 | 5784 | 0.9958 | 0.9976 | 0.9645 | 0.9036 | 0.9544 | 0.9385 | 0.9919 | 0.8921 |
|  | Polymorphic | 3131 | 737 | 0.9949 | 0.9988 | 0.9756 | 0.8657 | 0.9597 | 0.9564 | 0.9955 | 0.8073 |
| CLEAR-CLIP | EXACT | 0 | 0 | NA | NA | NA | NA | NA | NA | NA | NA |
|  | 5' | 13180 | 1225 | 0.9369 | 0.9937 | 0.9344 | 0.5393 | 0.8857 | 0.9836 | 0.8899 | 0.8400 |
|  | 3' | 8485 | 828 | 0.9418 | 0.9941 | 0.9439 | 0.5848 | 0.9018 | 0.9836 | 0.9072 | 0.8454 |
|  | Polymorphic | 4317 | 426 | 0.9167 | 0.9917 | 0.9136 | 0.4975 | 0.8533 | 0.9837 | 0.8529 | 0.8568 |

**Table S6:** Common pathways of the miRNA clusters in the CLASH 20% test data and CLEAR-CLIP data.

|  | Cluster | Number of common Targets | Common KEGG pathway |
| --- | --- | --- | --- |
| CLASH | hsa-miR-106a-5p, hsa-miR-20a-5p, hsa-miR-20b-5p, hsa-miR-26b-5p, hsa-miR-93-5p | 194 | Hepatitis B |
|  | hsa-miR-106a-5p, hsa-miR-17-5p, hsa-miR-20a-5p, hsa-miR-20b-5p, hsa-miR-26b-5p | 194 | Hepatitis B |
|  | hsa-miR-106a-5p, hsa-miR-17-5p, hsa-miR-196a-5p, hsa-miR-196b-5p, hsa-miR-20b-5p | 81 | Proteoglycans in cancer, TGF-beta signaling pathway, Hepatitis B, FoxO signaling pathway, Chronic myeloid leukemia |
|  | hsa-miR-106a-5p, hsa-miR-20a-5p, hsa-miR-20b-5p, hsa-miR-421 | 73 | TGF-beta signaling pathway, Lysine degradation |
|  | hsa-miR-181b-5p, hsa-miR-196a-5p, hsa-miR-196b-5p, hsa-miR-20b-5p | 33 | Proteoglycans in cancer, p53 signaling pathway |
|  | hsa-miR-181b-5p, hsa-miR-20a-5p, hsa-miR-20b-5p, hsa-miR-421 | 32 | p53 signaling pathway |
|  | hsa-let-7a-5p, hsa-let-7d-5p, hsa-miR-4516 | 4 | Lysine degradation |
|  | hsa-let-7d-5p, hsa-let-7g-5p, hsa-let-7i-5p | 807 | Cell cycle, Hippo signaling pathway, Viral carcinogenesis, Proteoglycans in cancer, Hepatitis B |
| CLEAR-CLIP | hsa-miR-106a-5p, hsa-miR-1299, hsa-miR-20a-5p, hsa-miR-20b-5p | 11 | Hepatitis B |
|  | hsa-miR-17-5p, hsa-miR-20a-5p, hsa-miR-20b-5p, hsa-miR-93-5p | 687 | Hepatitis B, Pathways in cancer, Proteoglycans in cancer, Chronic myeloid leukemia, FoxO signaling pathway |
|  | hsa-let-7a-5p, hsa-miR-1268a, hsa-miR-1277-5p | 1 | Pathways in cancer, Transcriptional misregulation in cancer |
|  | hsa-miR-320b, hsa-miR-320c, hsa-miR-320d | 513 | Hippo signaling pathway, Adherens junction |
|  | hsa-let-7a-5p, hsa-miR-1268a | 9 | Lysine degradation, Viral carcinogenesis, Bacterial invasion of epithelial cells, Chronic myeloid leukemia, Pathways in cancer, Transcriptional misregulation in cancer |
|  | hsa-miR-1268a, hsa-miR-1268b | 5 | Lysine degradation, Viral carcinogenesis, |
|  | hsa-miR-1277-5p, hsa-miR-138-5p | 27 | Pathways in cancer, Proteoglycans in cancer |

**Table S7:** Performance comparison between DMISO trained on top 5 pairs of miRNA/isomiR-mRNA target sites with external tools on CLASH 20% test, CLEAR-CLIP and miRTarBase datasets.

|  |  | Pos | Neg | AUROC | AUPR | F1 | MCC | Accuracy | Precision | Recall | Specificity |
| --- | --- | --- | --- | --- | --- | --- | --- | --- | --- | --- | --- |
| CLASH test | DMISO | 42561 | 25804 | 0.9942 | 0.9966 | 0.9667 | 0.9102 | 0.9579 | 0.9512 | 0.9827 | 0.9169 |
|  | miRanda | 42561 | 25804 | 0.6022 | 0.6987 | 0.3450 | 0.2929 | 0.5058 | 0.9869 | 0.2090 | 0.9954 |
|  | RNA22 | 42561 | 25804 | 0.5006 | 0.6229 | 0.0033 | 0.0155 | 0.3783 | 0.8353 | 0.0017 | 0.9995 |
|  | TargetScan | 42561 | 25804 | 0.5647 | 0.6695 | 0.2424 | 0.2195 | 0.4603 | 0.9612 | 0.1387 | 0.9908 |
|  | miRAW | 42561 | 25804 | 0.5497 | 0.6531 | 0.2678 | 0.1469 | 0.4543 | 0.8129 | 0.1603 | 0.9392 |
|  | miTAR | 42561 | 25804 | 0.5946 | 0.6779 | 0.4939 | 0.2018 | 0.5376 | 0.7753 | 0.3624 | 0.8268 |
| CLEAR-CLIP | DMISO | 45752 | 4388 | 0.9559 | 0.9954 | 0.9332 | 0.5699 | 0.8846 | 0.9894 | 0.8830 | 0.9011 |
|  | miRanda | 45752 | 4388 | 0.5285 | 0.9174 | 0.1414 | 0.0627 | 0.1554 | 0.9765 | 0.0762 | 0.9809 |
|  | RNA22 | 45752 | 4388 | 0.4996 | 0.9124 | 0.0010 | -0.0102 | 0.0879 | 0.7931 | 0.0005 | 0.9986 |
|  | TargetScan | 45752 | 4388 | 0.6124 | 0.9320 | 0.3967 | 0.1513 | 0.3118 | 0.9911 | 0.2480 | 0.9768 |
|  | miRAW | 45752 | 4388 | 0.5019 | 0.9128 | 0.1661 | 0.0038 | 0.1632 | 0.9158 | 0.0913 | 0.9125 |
|  | miTAR | 45752 | 4388 | 0.6756 | 0.9425 | 0.6294 | 0.2003 | 0.5012 | 0.9772 | 0.4642 | 0.8870 |
| miRTarBase | DMISO | 14144 | 0 | NA | NA | NA | NA | NA | NA | 0.8249 | NA |
|  | miRanda | 14144 | 0 | NA | NA | NA | NA | NA | NA | 0.7045 | NA |
|  | RNA22 | 14144 | 0 | NA | NA | NA | NA | NA | NA | 0.0199 | NA |
|  | TargetScan | 14144 | 0 | NA | NA | NA | NA | NA | NA | 0.7632 | NA |
|  | miRAW | 14144 | 0 | NA | NA | NA | NA | NA | NA | 0.6734 | NA |
|  | miTAR | 14144 | 0 | NA | NA | NA | NA | NA | NA | 0.0090 | NA |

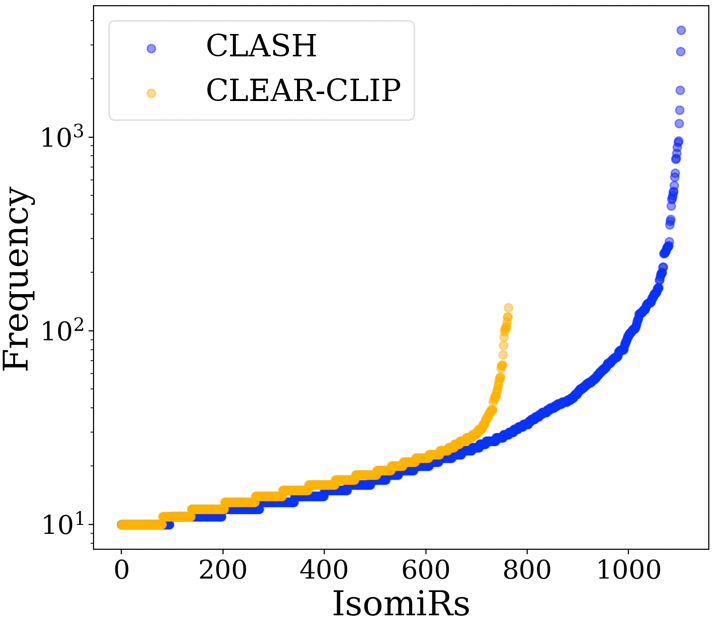

**Figure S1:** Frequencies of the 1,104 and 764 isomiRs in the CLASH and CLEAR-CLIP datasets respectively. The Y-axis is shown in log scale.

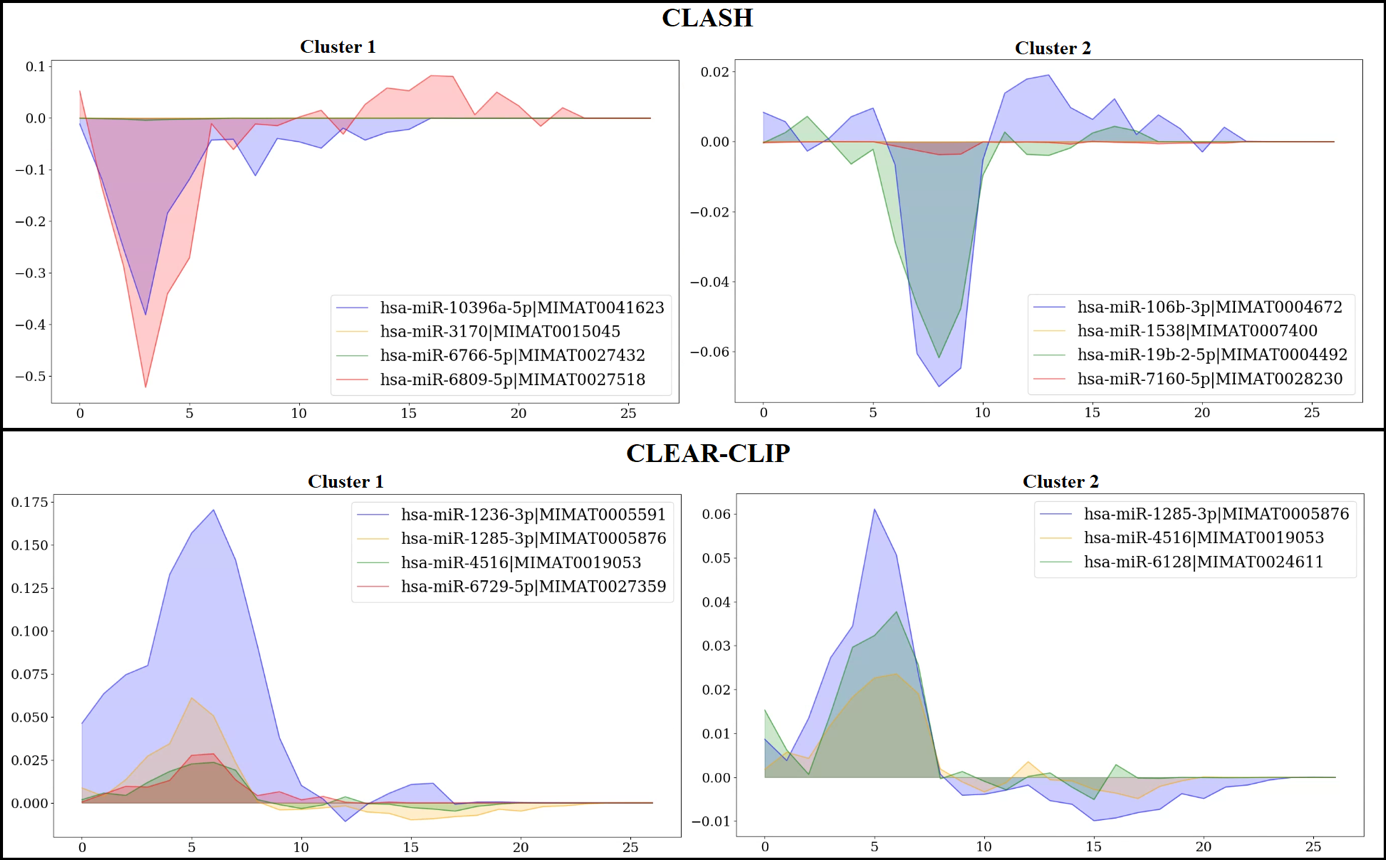

**Figure S2:** The cluster of miRNAs for which DMISO reacted similarly to the position-wise changes. The top 2 clusters of miRNAs are shown for the two datasets. The X-axis represents the miRNA positions and Y-axis represents the changes in DMISO’s prediction based on the changes in corresponding miRNA positions.
